# The Biological Basis of the Experience of Constant Colour Categories

**DOI:** 10.1101/488379

**Authors:** Semir Zeki, Alexander Javier, Dimitris Mylonas

**Affiliations:** Laboratory of Neurobiology, University College London

**Keywords:** colour, vision, biological categories, constant colour categories

## Abstract

In the work reported here, we used the Land Colour Mondrian experiments to test the degree to which subjects vary in their perception of colour categories. Twenty subjects of different ethnic and cultural backgrounds, for all but one of whom English was not the primary language, viewed 8 patches of different colour in two Mondrian displays; each patch, when viewed, was made to reflect identical ratios of long-, middle- and short-wave light. Subjects were asked to match the colour of the viewed patch with that of the Munsell chip coming closest in colour to that of the viewed patch, without using language. Overall, there was no variability in categorizing colours as red, yellow, brown and green but a small variability in categorizing them as purple, blue, orange and turquoise. In terms of hue, we found significantly less variability in matching ‘warm’ hues than in ‘cool’ ones. We interpret the lack of significant variability between subjects in the matches made as a pointer to similar computational mechanisms being employed in different subjects to perceive colours, thus permitting subjects to assume that their categorization of colours has universal agreement and assent.

## Introduction

It is trite neurobiology to say that one of the primordial functions of the brain is to acquire knowledge. Yet it raises a fundamental issue of huge importance, namely the extent to which the knowledge acquired through brain mechanisms by one individual is identical to that acquired by another or others, thus allowing the acquiring individual to assume reasonably that there is universal assent to the knowledge and the experience acquired by him or her. The question then resolves itself around asking what the conditions are which enable all individuals, irrespective of their ethnic, cultural or educational status, to share the same experience and knowledge under the same conditions.

There has been much philosophical debate about this subject, which we do not delve into here in any detail. Rather, accepting Immanuel Kant’s [1] statement that “perceptions without concepts are blind”, we work on the assumption that all sensory inputs are interfaced through brain concepts, of which, we believe, there are two kinds: inherited and acquired (synthetic) [2]. The former results in knowledge which, to a greater or lesser extent, is similar in all individuals, thus making it easy for one individual to assume that others would share the same or nearly identical experience under similar conditions; the latter, by contrast, lead to experiences and knowledge that can differ profoundly between individuals, even if experienced under similar conditions, making it unsafe for an individual to assume universal assent to his or her experience. In more general Bayesian terms, we consider the former, inherited, concepts to generate biological priors (*β priors*) which are similar, and resistant to modification through experience, in all individuals, of which one example is that of faces or human bodies; in the other category are artefactual priors (*α priors*) such as those of cars or other manufactured goods, which differ between individuals, are much more dependent upon culture and learning and are modifiable through experience [3].

The most extreme example of a biological *β prior* generated by interfacing incoming visual signals with an inherited, biological, mechanism is that of colour vision. It is common knowledge that the colour category to which objects and surfaces belong do not change with fairly wide ranging changes in the wavelength-energy composition of the light in which they are viewed [4] [6], a phenomenon generally referred to as colour constancy. We prefer to use the term “constant colour category”, because the hue (shade) of colour of a given surface or object does change with changes in the wavelength-energy composition of the light in which it is viewed, even if the colour category to which it belongs does not [7].

The distinction between *colour* and *colour category* is important for the experiments described in this paper, in which we set out to learn the extent of variability in the experience of colour categories when individuals of different ethnic and cultural backgrounds view the same stimuli under the same conditions. Our present paper is a further step in a broader experimental one, which aims to address the degree of variability produced in individuals when their experience is the product of interfacing the incoming visual signals with inherited and acquired brain concepts. The first study addressed the question of the biological basis of mathematical beauty [8].

No one has determined the precise concept, in neural terms, which the brain uses to generate colours. But the ratio-taking formulations produced by Edwin Land and his colleagues [4] [5] [6] are perhaps the most valid currently available and the easiest to use, given their mathematical precision and the predictability of the results produced through them. The exact concept is in any case not critical for the work reported here but the experiments we have used are, and these are based on Land’s. Our aim was very simple and directed toward learning the extent to which different individuals’ categorization of colour is the same. Our hypothesis was that there would be negligible difference between individuals in this categorization. This may seem obvious but was important to demonstrate formally.

Berlin and Kay [9] proposed a total universal inventory of eleven basic colour categories (corresponding to the English terms: white, black, red, yellow, green and blue, grey, purple, brown, orange and pink) defined by a combination of multiple linguistic and psychological criteria. Such criteria have been strongly criticized as being not equally applicable across languages [10][11]. Others have identified basic colour categories on more objective behavioural criteria such as frequency, response time and consistency of colour naming responses [12] [13] [14]. Recently colour categories have been studied by measuring the communication effectiveness of colour names using information theoretic analysis [15] [16]. Because these approaches have been based on the use of language and because variations in categorizations are traceable to language, we opted for a different approach. We investigated whether subjects of different linguistic and ethnic backgrounds categorize different colours in a similar way, when light reflected from patches of different colour have the same wavelength-energy composition. We did so by asking subjects to match the experienced colours of the viewed patches with that of a standard set of Munsell chips, without the use of language.

## Material & Methods

### Subjects

Twenty subjects, of whom 10 were females, took part in the experiment; their mean age was 24.1 years; Standard Deviation (SD) was 6.7. They were recruited through advertisements at University College London (UCL), were over 18 years of age and had normal or corrected to normal vision. They were all tested with Ishihara plates (Ishihara, 1988) for colour vision abnormalities and none was found to be deficient. No subject reported any neurological or psychological disorder, all gave informed consent and the experiment was approved by the UCL Ethics Committee (12327/001). Subjects came from the following countries: Cyprus, Thailand, Turkey, Lebanon, France, Ghana, China, Brazil, India and Spain; to all but one English was a second language.

### Task

Subjects had to match the colour of patches in a Land Colour Mondrian display under specific conditions of illumination (see below) with coloured Munsell chips viewed under specific illumination conditions, thus obviating the use of language.

### The Mondrian displays

As in Land’s original experiments, we used Land Colour Mondrian displays, which were placed at a distance of 2 metres from the observers. These displays consist of an assembly of squares and rectangles so arranged as to form an abstract scene with no recognizable shapes or objects, besides rectangles and squares. This controls for any effects due to memory and learning of what colours objects should have. To avoid specular reflectance, we used matt Colour Aid papers, which reflect a constant amount of light in all directions. No patch was surrounded by another patch or patches of a single colour, thus avoiding induction effects.

We constructed and used two Mondrian displays, and subjects had to match the colours in each (see Figure 1) to Munsell chips. Eight test patches were selected in each display, based on their proximity in CIELAB colour space (mean ΔE00 = 5.5, SD = 3.4, see Figure S1) to the centroids of the proposed basic colour categories in English: Blue, Brown, Green, Orange, Purple, Red, Turquoise and Yellow, as identified by an online colour naming experiment [14]. In the following description, each of the test patches will be referred to by the colour names given above (Figure 1) although in the experiments subjects did not use language but merely matched the patch to the Munsell chips. Both Mondrian displays included the same eight test colour patches but in different configurations; in both, each patch subtended 8.25° x 6° and the surrounding patches extended more than 10° in all directions.

**Figure 1.**
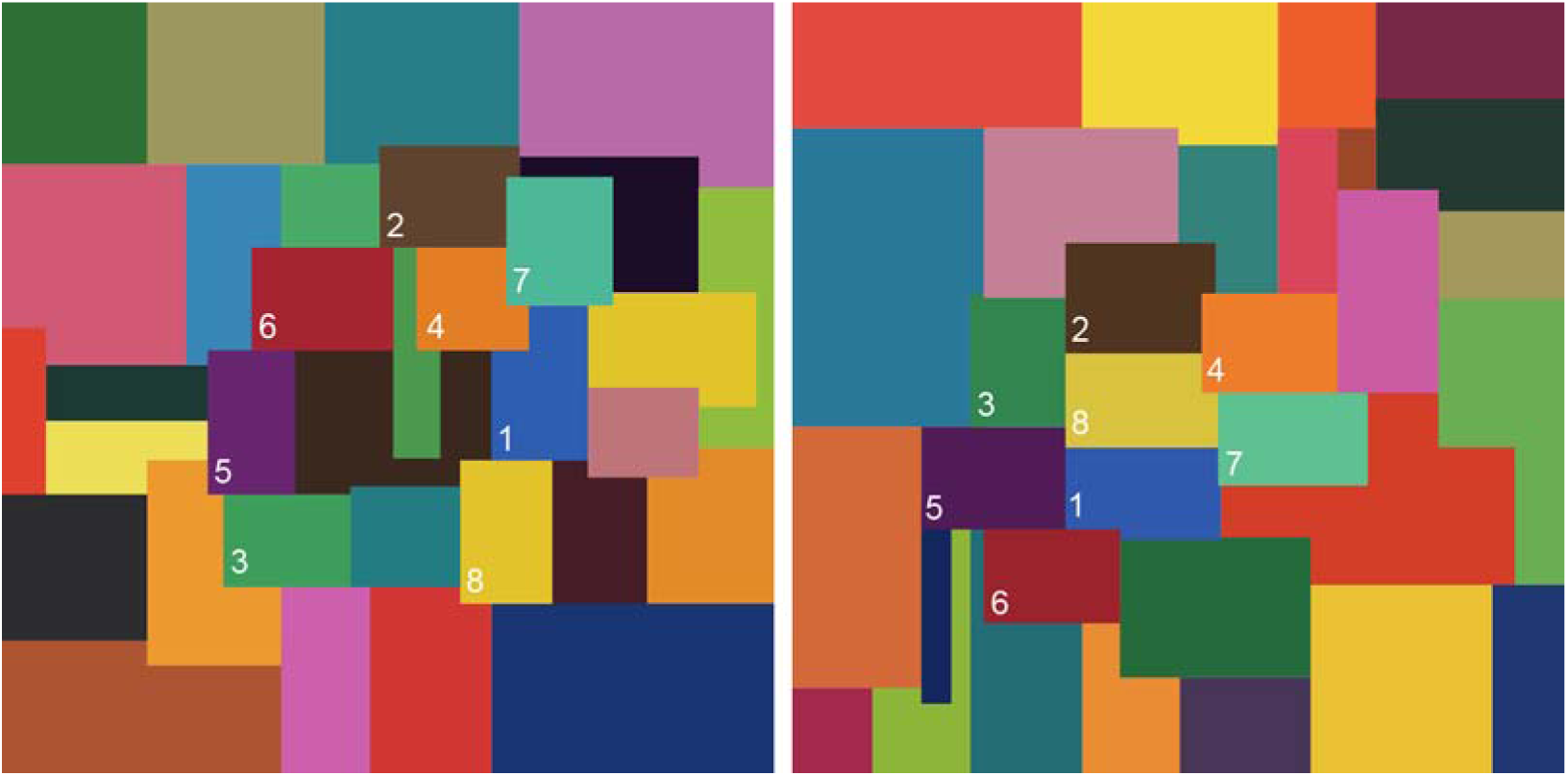
Appearance of Mondrian display 1 (left) and Mondrian display 2 (right) under daylight viewing conditions. 1=Blue, 2=Brown, 3=Green, 4=Orange, 5=Purple, 6=Red, 7=Turquoise, 8=Yellow.

The Mondrian displays were illuminated by three carousel projectors (Kodak Ektagraphic B-2AR), equipped with ELH 120V 300W bulbs, rheostats and three gelatine filters passing long-, middle-, and short-wave light, respectively; the filters had been specially manufactured for Zeki’s experiments by Edwin Land [17]. The long-wave filter transmitted light in the range of 592nm to the long end of the visible spectrum with a peak transmittance greater than 660nm. The transmittance of the middle-wave filter was in the range 492-580nm (peak 528nm) while the short-wave filter transmitted light in the range 386-493nm (peak 432 nm) with a secondary peak at 700nm. Each projector was equipped with a separate rheostat and shutter, thus enabling the intensity of light coming from each to be adjusted separately.

For each test stimulus, we adjusted the amount of long, middle and short wave light of the three carousel projectors so that each patch, when judged for its colour, reflected a nearly constant ratio of 60% long-, 20% middle- and 20% short- wave light. The energies reflected from each patch were measured in milliwatts per steradian per square meter (mW.Sr^-1^.m^-2^) separately for each projector using a PR-670 tele-spectroradiometer (Figure 2). We also report the stimulus specifications for each test patch in 10° relative cone excitation units [18] in Supplementary Figure S2. The consistency of the ratios was checked before each experimental session.

**Figure 2.**
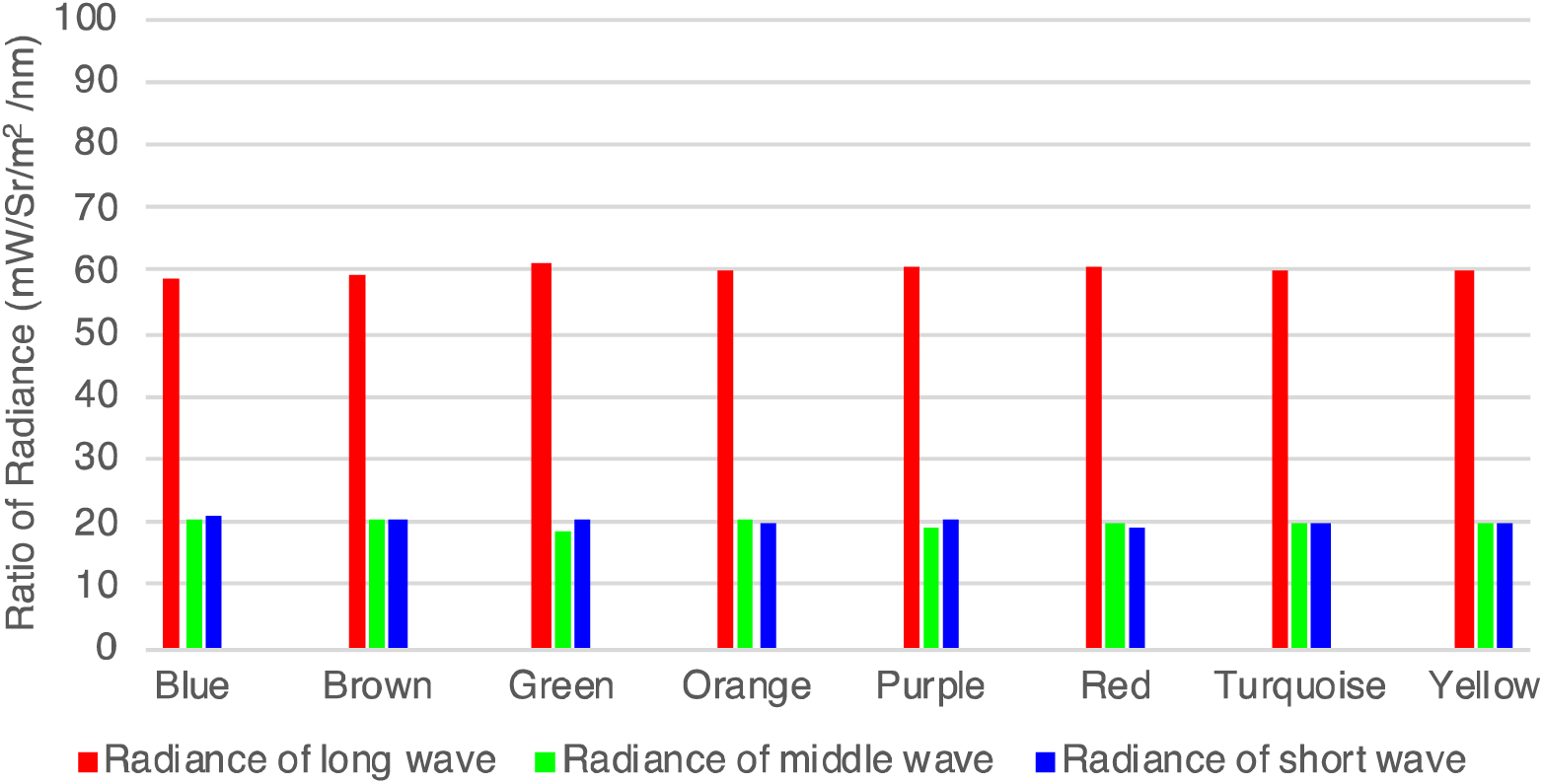
Ratio of radiances (mW/Sr^-1^/m^-2^) for long-, middle- and short wave light reflected from each test patch in the Mondrian displays.

### The Munsell Chips

Under the illumination conditions specified above, subjects were asked to match the colour of the nominated patches with one of the 44 colour chips from the Munsell Book of Color (Glossy Collection, M40115). The 40 hues of the Munsell set were selected to have the maximum available chroma (saturation) and variable Value (lightness) levels. For the yellow to red hues we also added four darker stimuli (Munsell 2.5YR to 10YR) because there were no brown or yellow chips at the same Value level (see Figure 3 and Supplementary Table S1 for the full specifications of the chips). The order of the chips was randomised and displayed on an annulus at a constant eccentricity of 10° visual angle; they were presented against a mid-neutral grey surround inside a viewing booth illuminated by two GrafiLite daylight simulators (CIE 1931 x=0.327, y=0.339).

**Figure 3.**
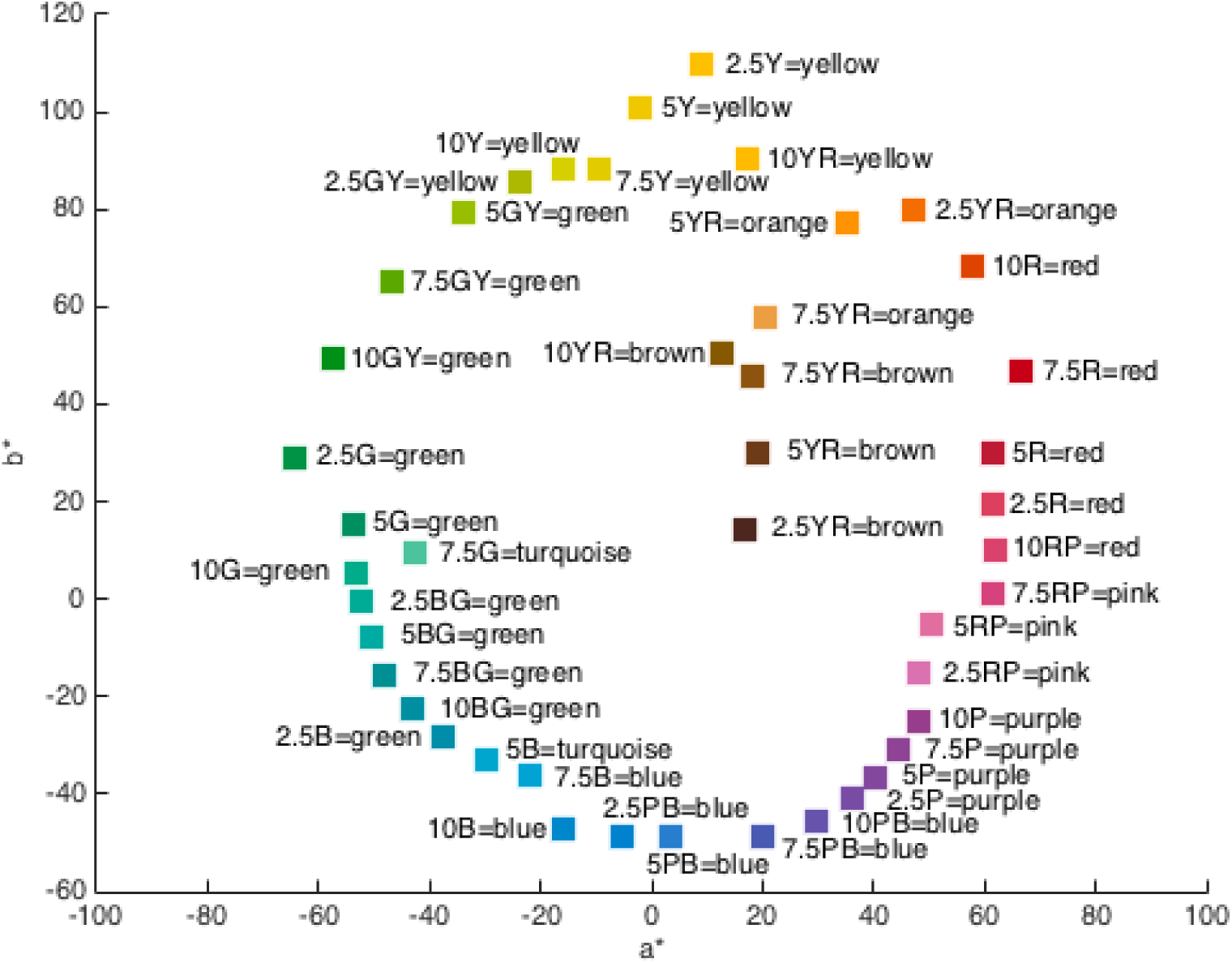
Munsell chips shown as though viewed from above (a* b* plane of CIELAB space) segregated according to colour name.

### Colour Matching Procedure

Participants had to match the colour of the eight test patches in each Mondrian display with the Munsell chip that, to them, was closest in colour to the patch under examination. The viewing booth with the Munsell chips was placed on a desk at a distance of 60cm from the observers. After adjusting the rheostats of the projectors to make each patch reflect the (same) amounts of long-, middle-, and short-wave light (given above), all three projectors were switched on to illuminate the entire Mondrian display, while the two daylight simulators were switched on to illuminate the 44 Munsell chips. Participants performed the successive colour matching tasks without time limit but, in practice, each trial took less than 1 minute. The procedure was repeated twice for each test patch using the two different Mondrian displays of Figure 1 to measure the reliability of the responses, thus giving a total of 16 trials per subject. Any remaining light sources in the experimental room were eliminated.

### Classification of hues into lexical colour categories

The exact hue or shade of a coloured surface varies under different conditions of illumination while its colour category remains constant [7]. We classified the hues of the chips chosen by the subjects into different lexical categories, based on a probabilistic colour naming model [19]. In Figure 3, we show the Munsell chips of our comparison stimuli assigned to the most likely colour names.

## Results

The means and standard deviations of the matches (between the Munsell chips and each of the eight test patches) for all subjects for both Mondrian displays are shown in Figure 4.

**Figure 4.**
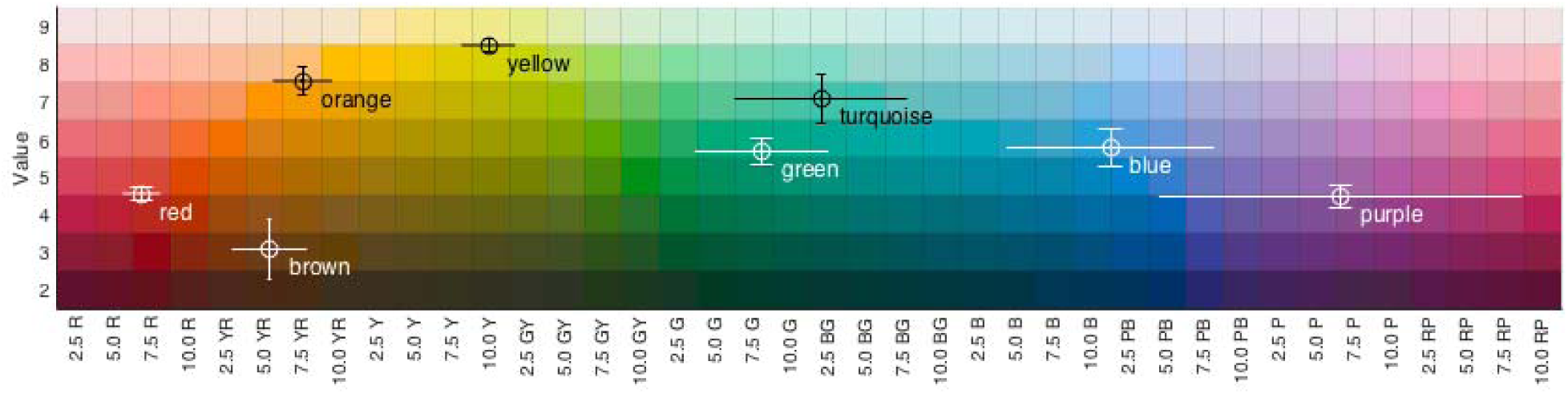
Variability of colour matching responses for each of the eight test patches for both Mondrian displays. Rings denote the mean and the error bars the standard deviation. The horizontal lines show the variability in the hue dimension and the vertical lines the variance in the Value dimension.

The patches with the lowest hue variance in their matched Munsell chips were red followed by yellow, orange and brown. The highest variability was observed for purple followed by blue, turquoise and green. Thus, the variability in colour matching responses is lower for reddish than bluish colours (r = 0.94, n = 8, p < 0.0005); this reflects the smaller perceptual extents of categories in the warm region (in terms of steps leading to a change in hue), than in the cool region of colour space [14].

Figure 5 shows the mean and standard deviation of the responses for each Mondrian display separately. A comparison of the means obtained for the different patches in the two Mondrian displays produced a good agreement (mean CIE ΔE 2000=2.00), with the largest differences being observed for the brown and purple patches (ΔE00=4.58 and ΔE00=2.23, respectively) and the smallest for the yellow patch (ΔE00=1.08). The variance for the purple patch was also wider for display 1 than for display 2, presumably because the former purple patch was surrounded by darker neighbouring patches (see patches 5 in Figure 1).

**Figure 5.**
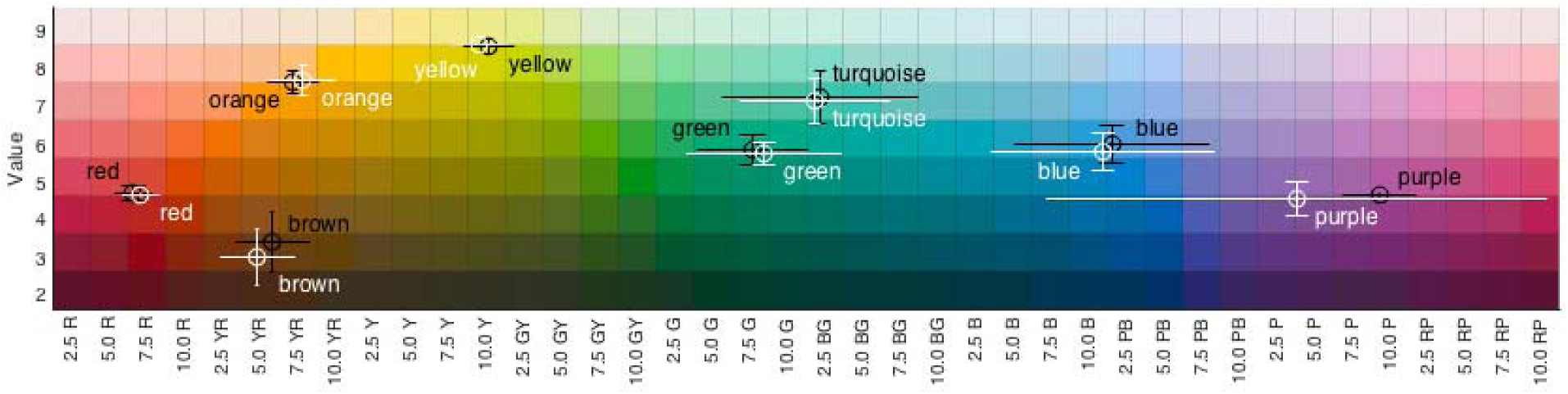
Variability of colour matching responses for each of the eight test patches for Mondrian display 1 (white) and Mondrian display 2 (black). Conventions as in Figure 4.

The above description applies to hues of the Munsell chips; we were in fact more interested in the variability of colour categories, because it is the colour category rather than the hue that remains constant [7]. In Figure 6, we convert the matches given in terms of the Munsell chips into lexical categories. There is no variability for matching the red, yellow, brown and green patches to their corresponding Munsell categories and high consistency for purple, orange, blue and turquoise. Despite the larger variability for the turquoise compared to the other patches, the allocation with regard to the same colour category in terms of Munsell chips was significant (*χ*^2^ 4.05, *p= 0.04, using Yate’s correction*), from which it follows that all the above matches were also significant.

**Figure 6.**
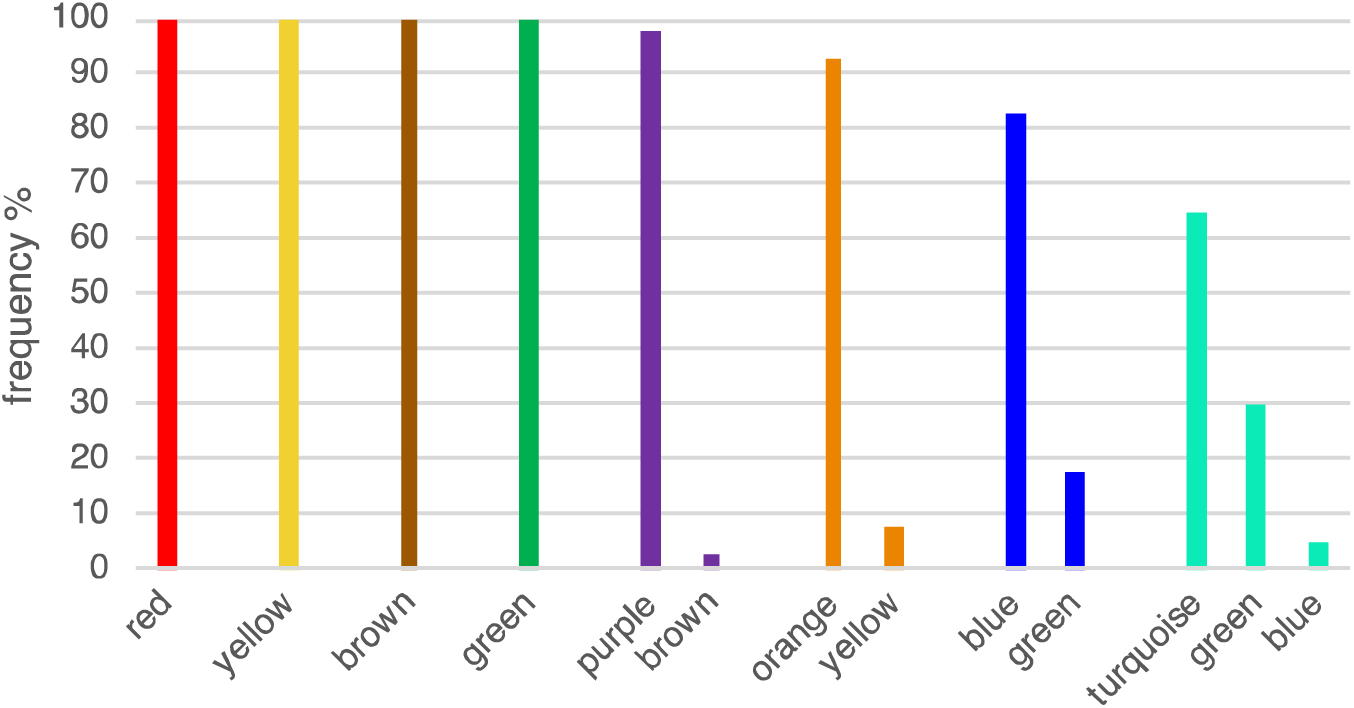
Frequency of corresponding colour categories of Munsell chips selected as the best match for each test patch in both Mondrians. Colours denote the colour of the test patch and the labels show the corresponding colour names for the Munsell.

Our results are summarized in Figure 7, which shows that of a total of 320 responses, 92% allocated the test patches of the Mondrian displays to the Munsell chips belonging to the same category while 8% were matched to chips belonging to other, but closely neighbouring, colour categories (*χ*^2^ = 448.9, *p < 0.0001*).

**Figure 7.**
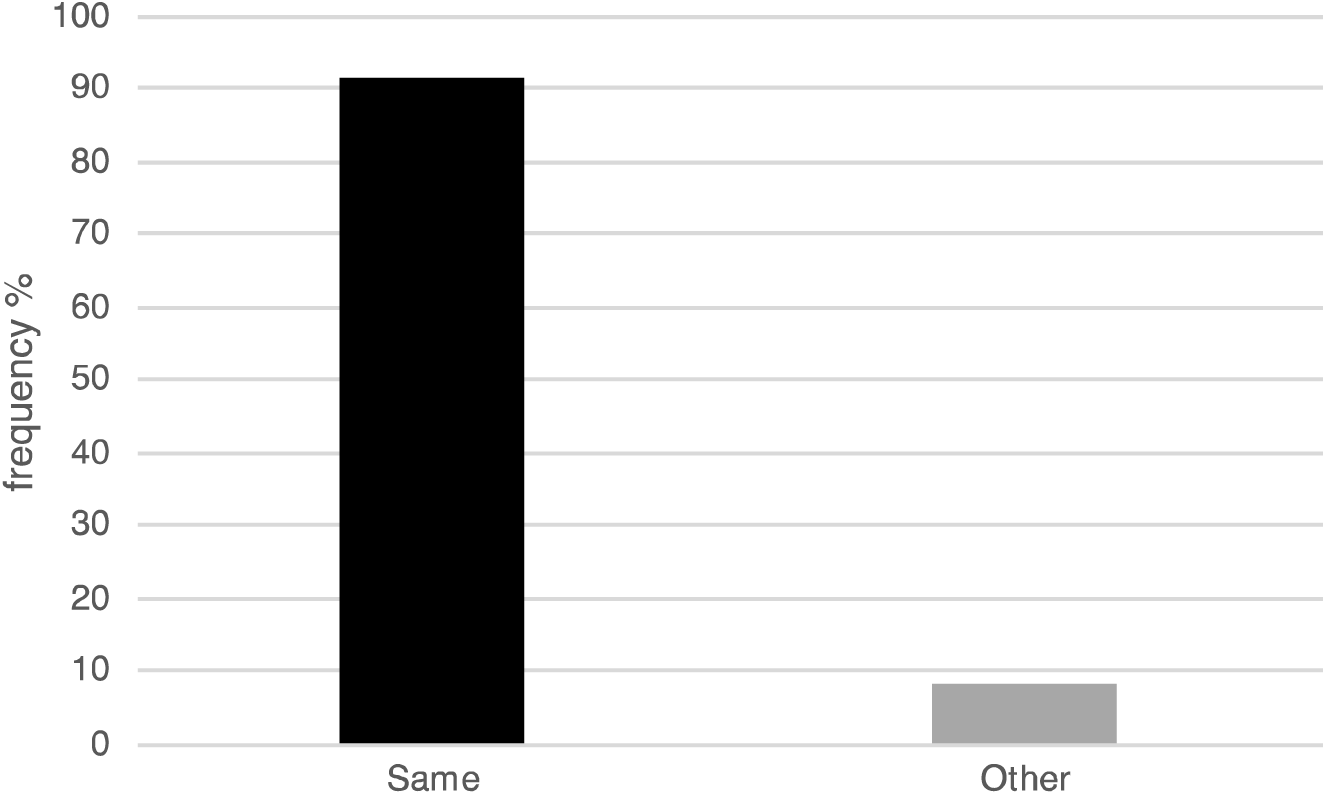
Proportions of responses belonging to the same or other category for both Mondrian displays.

## Discussion

The experiments reported here constitute part of a series in which we explore judgments that can reasonably be accounted for by supposing that they are based on biologically inherited concepts or mechanisms and are thus distinct from post-natally acquired ones [2]. Certain characteristics facilitate the categorization of experiences or judgments as being based predominantly or even exclusively on biologically inherited concepts. Prominent among these is a lesser variability between subjects, even those belonging to different races and cultures, when making judgments based on inherited concepts [3]. The consequence of this more restricted variability is that the individual making a judgment based on inherited concepts is more entitled to assume that his or her judgment has universal validity and assent. This has so far been found to be true for aesthetic judgments of portraits and landscapes [20] as well as mathematical formulae experienced as beautiful [8], all of which we consider to belong to the biological category. Aesthetic judgments based on such concepts are characterized by lesser variability in judgment ratings compared to aesthetic judgments of man-made artefacts such as buildings (which are more likely to be interfaced through synthetic concepts). In the work reported here, we extend this approach to colour vision.

### Colour Categorization is dictated by inherited programmes or concepts

Colour is perhaps the most extreme example of an experience that is dictated by an inherited brain concept. We refer to this concept, based largely on the work of Edwin Land and his colleagues, as a ratio-taking concept (although one could equally refer to it as a brain mechanism or programme). Specifically, the concept is one in which light of any waveband reflected from a surface is compared with light of the same waveband reflected from surrounding surfaces, and a ratio between the two taken. Although Land supposed that this is done three times, for long-, middle- and short-wave light, it is equally possible that it is done many times for lights of many different wavebands. The net result of these operations is that colour perception becomes largely independent of the continuous fluctuations in the wavelength-energy composition of the light reflected from a surface, thus leading to a perceptual stabilization of colours. It is common to suppose and write of the result of such a stabilization as colour constancy, by which is meant that the colours generated are constant and largely independent of the precise wavelength-energy composition of the light reflected off them. We believe, however, that there is a better way of describing the end result, because what does not change as a result of this ratio-taking operation is in fact the colour category, not the hue (or shade of colour), which changes when surfaces are viewed in different illuminants [7]. Consistent with this belief, what we have shown here, in summary, is that colour categories remain stable and that such variation as there is, is rather in the hues within these constant colour categories that change.

We undertook this study in the belief that the result of ratio-taking operations are similar in all humans. The consequence is that the results will also be similar in all humans, with little variability in the ascription of colours to given categories. The variability in matching the colour of the patches with chips belonging to different Munsell categories was indeed very limited, especially for red, yellow and orange (warm colours) while it was broadest for purple and green (cool colours). The small variance in the red and yellow matches reflects the fact that, in terms of their extent measured as steps leading to a change in hue, they are indeed smaller categories than green or purple, which have a larger number of hue steps and hence a higher variability [14].

This difference in the width between warm and cool colours, in terms of hue steps needed to change to another colour category, has been observed before. The work of Gibson and his colleagues [16] showed that warm colour categories are more salient than cool ones and Danilova and Mollon [21], in their colour discrimination studies, showed a border between warm and cool categories, which we interpret as signifying that warm colours are narrower in terms of hue steps than cool ones. Although our results correspond to theirs in drawing a distinction between warm and cool colours, we do not interpret our results, as they do, in terms either of post-receptoral channels [21] or of communication needs [16]. For us, the task was strictly a computational perceptual task and reflects the constancy of colour categories, reached by a computational brain process that is independent of learning, memory, environmental and social factors. And, while acknowledging that linguistic criteria may be of importance in classifying colours in terms of language, we believe that the classification according to colour categories is not dependent upon language and experience. Supporting evidence for this comes from the ability of children and monkeys to categorize colours much like adult humans [22][23][24].

That the ratio taking operation we describe above, or some other computational paradigm very similar to it, should lead to constant colour categories raises interesting questions from a Bayesian point of view [3]. Specifically, a colour category can never become a posterior; it is always a prior. This is because, no matter what the wavelength energy composition of the light reflected from, say, a green patch, it will always belong to the green category. Only the hues within that patch can become posteriors which can then act as priors for the generation of other (posterior) hues but ones which belong to the same colour category.

### Conclusion

In summary, there was a trivial variability in assigning colours to different categories by subjects of different ethnic and cultural origins. This is a pointer to an important principle of the organization of the sensory brain, at least in terms of colour vision, namely that there is a very significant similarity in the inherited computational mechanisms for generating colour categories in all humans.

## Data Accessibility

Psychophysical data, colorimetric data and code: Dryad doi:10.5061/dryad.hd593r6 [25]

## Authors’ contributions

SZ conceived and designed the study and authored the manuscript. AJ carried out the colorimetric measurements and collected the psychophysical data. DM coordinated the study, carried out the colour stimulus specifications and statistical analyses, and helped draft the manuscript. All authors gave approval for publication.

## Data Accessibility

Psychophysical data, colorimetric data and code: Dryad doi:10.5061/dryad.hd593r6 [25]

## Supplementary Material

**Figure S1.**
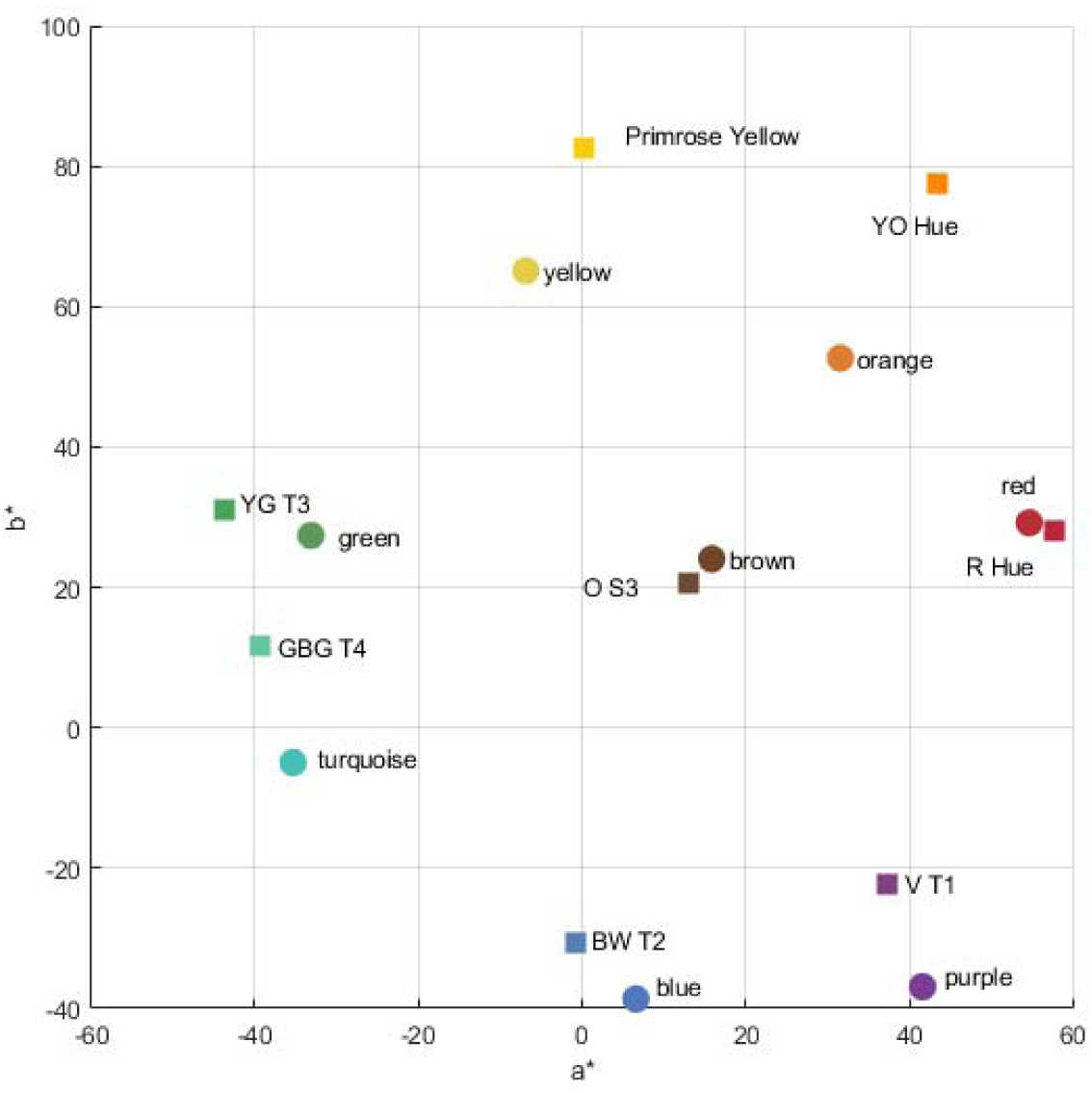
Centroids of colour names (circles) reported by Mylonas & MacDonald (2016) and test patches of Color-aid papers (squares) in CIELAB.

**Figure S2.**
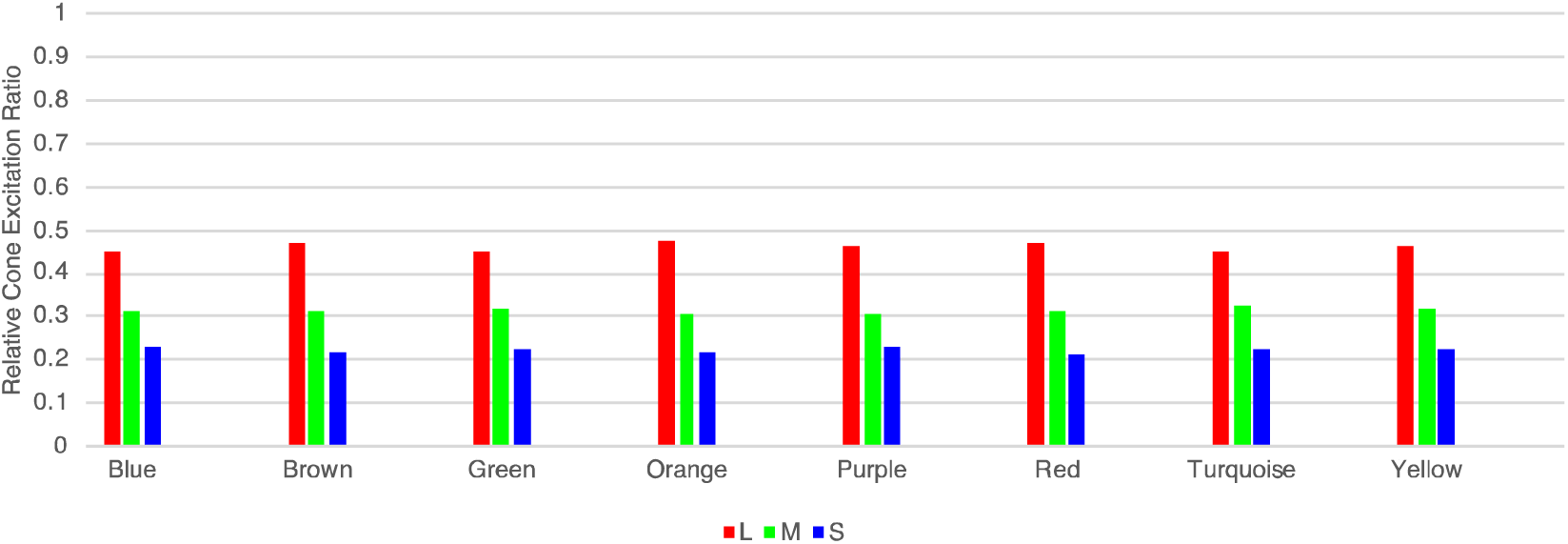
Long (L), Medium (M) and Short (S) cone excitation ratios [18] in 10° for each of the eight test stimuli of the Mondrian displays.

**Table S1.**
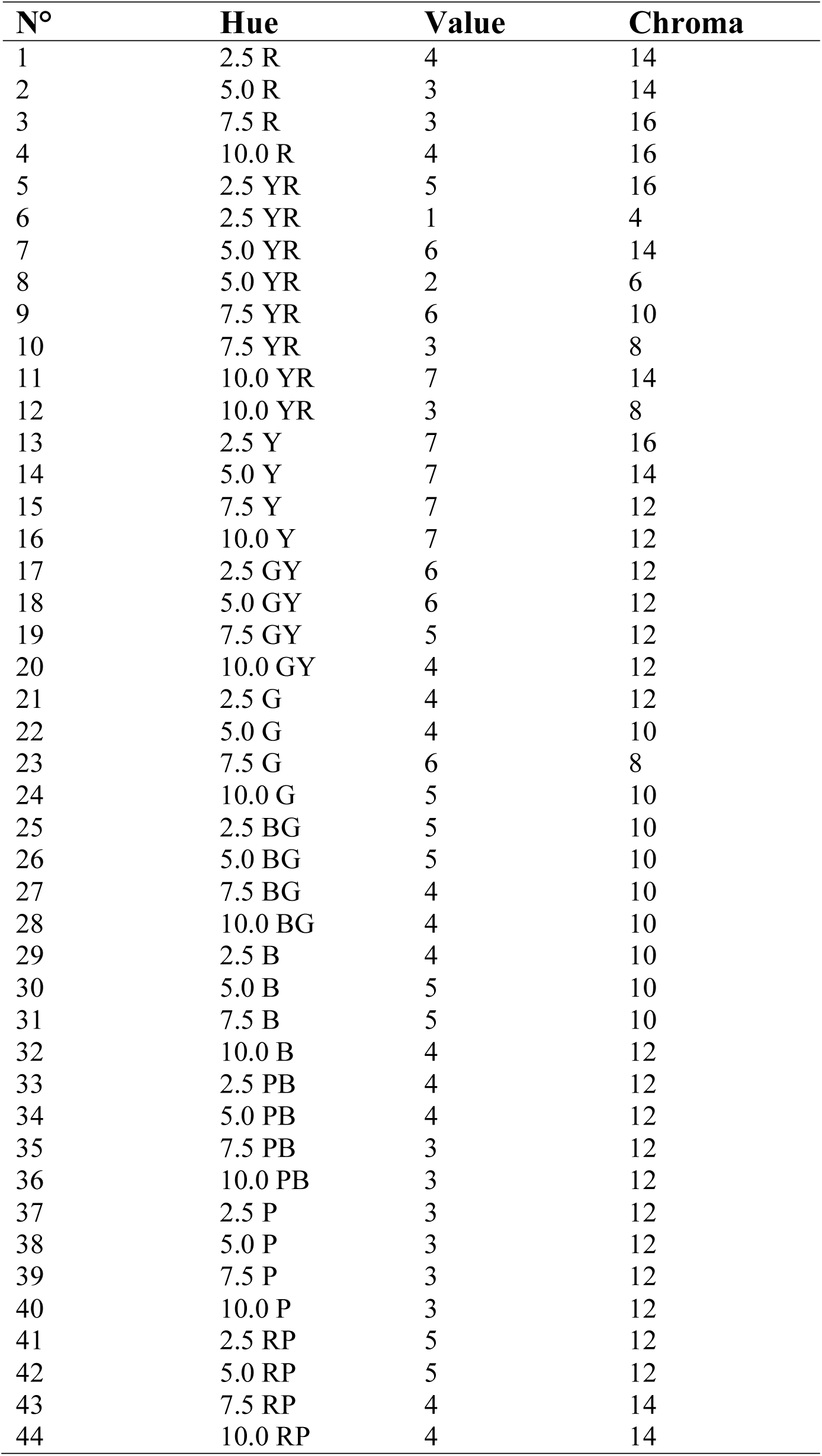
Munsell notation of comparison stimuli.

